# Genome scale mutational analysis of *Geobacter sulfurreducens* reveals distinct molecular mechanisms for respiration of poised electrodes vs. Fe(III) oxides

**DOI:** 10.1101/084228

**Authors:** Chi Ho Chan, Caleb E. Levara, Fernanda Jiménez-Oteroa, Daniel R. Bond

## Abstract

*Geobacter sulfurreducens* generates electricity by coupling intracellular oxidation of organic acids with electron transfer to the cell exterior, while maintaining a conductive connection to electrode surfaces. This unique ability has been attributed to the bacterium’s capacity to also respire extracellular terminal electron acceptors that require contact, such as insoluble metal oxides. To expand the molecular understanding of electricity generation mechanisms, we constructed *Geobacter sulfurreducens* transposon mutant (Tn-Seq) libraries for growth with soluble fumarate or an electrode surface as the electron acceptor. Mutant libraries with over 33,000 unique transposon insertions and an average of 9 transposon insertions per kb allowed identification of 1,214 genomic features essential for growth with fumarate, including over 270 genes with one or more functional homologs that could not be resolved by previous annotation or *in silico* modeling. Tn-Seq analysis of electrode-grown cells identified mutations in over 50 genes encoding cytochromes, processing systems for proline-rich proteins, sensory systems, extracellular structures, polysaccharides, metabolic enzymes and hypothetical proteins that caused at least a 50% reduction in apparent growth rate. Scarless deletion mutants of genes identified via Tn-Seq revealed a new putative *c*-type cytochrome conduit complex (*extABCD*) essential for growth with electrodes, which was not required for Fe(III)-oxide reduction. In addition, mutants lacking components of a putative methyl-accepting chemotaxis/cyclic dinucleotide sensing network (*esnABCD*) were defective in electrode growth, but grew normally with Fe(III)-oxides. These results suggest that *G. sulfurreducens* possesses distinct mechanisms for recognition, colonization, and reduction of electrodes compared to other environmental electron acceptors.

## Importance

Many metal-reducing organisms can also generate electricity at anodes. Because metal oxide electron acceptors are insoluble, one hypothesis is that cells sense and reduce metal particles using the same molecular mechanisms used to form biofilms on electrodes and produce electricity. However, by simultaneously comparing thousands of *Geobacter sulfurreducens* transposon mutants undergoing electrode-dependent respiration, we discovered new cytochromes and chemosensory proteins essential for growth with electrodes that are not required for metal respiration. This supports an emerging hypothesis where *G. sulfurreducens* recognizes surfaces and forms conductive biofilms using sensing and electron transfer pathways distinct from those used for growth with metal oxides. These findings provide a molecular explanation for studies that correlate electricity generation on electrode surfaces with direct interspecies electron transfer rather than metal reduction by *Geobacter* species, and reveal many previously unrecognized targets for improving and engineering this biotechnologically useful capability in other organisms.

## Introduction

While most electron acceptors are soluble and easily reduced by inner membrane respiratory proteins, some electron acceptors are insoluble or lie beyond the cell surface. Bacteria able to catalyze ‘extracellular electron transfer’ or ‘extracellular respiration’ contain redox proteins and attachment mechanisms able to create electrical connections between membranes and these external substrates. Multiple modes of extracellular respiration are known, including reduction of Fe(III) and Mn(IV) oxides (1, 2), electricity production at electrode surfaces (3, 4), and formation of between-cell conductive networks enabling syntrophic growth with electron-consuming methanogens (5, 6).

The facultative anaerobe *Shewanella oneidensis* uses a single pathway comprised of the CymA inner membrane cytochrome and the MtrABC porin-multiheme *c-*type cytochrome complex to deliver electrons to all tested external metals and electrodes (7, 8). In contrast, there is evidence for multiple levels of complexity in the respiratory chain of the anaerobe *Geobacter sulfurreducens. G. sulfurreducens* requires two different inner membrane *c*-type cytochromes, ImcH and CbcL, depending on the reduction potential of the extracellular acceptor (9, 10). Up to five homologs of the PpcA periplasmic tri-heme *c*-type cytochrome could be involved in assisting electron transfer across the periplasm (11). To cross the outer membrane, one electron transfer ‘conduit’ complex consisting of the OmaB multiheme *c-*type cytochrome, OmbB porin-like protein, and OmcB multiheme lipoprotein *c-*type cytochrome is essential for growth with some metals (12, 13), but the genome encodes at least five other putative porin-cytochrome complexes with unknown functions. At the outer surface, *Geobacter* must build a conductive interface to reach metal particles, electrode surfaces, or partner bacteria. Simple extracellular polysaccharides influence biofilm interactions (14), while a combination of conductive Type IV pili and multiheme *c-*type cytochromes such as OmcS, OmcE, OmcZ and PgcA are implicated in electron transfer beyond the cell membrane, depending on growth conditions (12, 14–16).

Due to the ease of screening mutants in liquid medium, compared to growth on electrodes in electrochemical reactors, most components of the *Geobacter* electron transfer pathway are identified using metals as proxies for other extracellular acceptors (14). Transposon-based mutagenesis using metals has revealed crucial cytochromes and attachment strategies, but high-throughput colorimetric screens are designed to identify mutants with growth rates near zero. Unfortunately, as multiple overlapping electron transfer pathways exist and many electron transfer proteins have functional homologs, single mutations typically decrease rather than eliminate growth of *G. sulfurreducens* (17, 18). This respiratory complexity, combined with the bottleneck of studying electron transfer in electrode biofilms, severely limits the pace of discovery when studying the molecular basis of complex phenotypes such as electricity production.

Transposon-insertion sequencing, also known as Tn-Seq, is able to quantify the abundance of every mutant containing a transposon insertion within a library, without individual mutant isolation (19). Using Tn-Seq, it is possible to measure the effect every mutation has on growth rate, by comparing an insertion’s abundance before and after a known number of generations. This method rapidly assessed gene essentiality under aerobic versus anaerobic growth in *Shewanella oneidensis*, revealed genes essential to different respiration strategies in *Rhodopseudomonas palustris*, and uncovered the essential gene set in the archaeon *Methanococcus maripaludis* (20–22). For this report, we constructed the first *G. sulfurreducens* Tn-Seq library, and used it to define over 1,200 genes essential for growth in minimal medium using a soluble electron acceptor. By then cultivating this same mutant library on electrode surfaces, a subset of over 50 additional genes that caused at least a 50% decrease in growth rate was revealed. Surprisingly, mutants lacking these new key cytochromes or chemosensory genes were severely impaired in growth with electrodes, but remained fully able to respire using other extracellular metal acceptors. These results suggest that separate biochemical mechanisms are used for electron transfer to electrodes and to environmental metals.

## Results and discussion

### Tn-Seq in *G. sulfurreducens.*

*G. sulfurreducens* transposon libraries were generated using a *mariner*-based transposon with directional type IIS MmeI recognition sites engineered on both ends of the transposon insertion sequence (22, 23). As MmeI cleaves chromosomal DNA 20 base pairs from its recognition site, adapter ligation and amplification allows sequencing to identify most insertions and quantify each mutant’s abundance (24). After three separate transposon libraries were constructed using fumarate as the electron acceptor, approximately 50,000 individual colonies were recovered per library, and sequencing revealed between 30,000 and 33,000 independent insertions in each library with no obvious hotspots or biases in coverage. The library representing the deepest coverage was used as the source for all experiments reported in this work.

When DNA from two separate cultures within the same library was extracted, digested with MmeI, ligated to Illumina adaptors, amplified, and sequenced, 33,257 and 33,343 unique insertions were identified in the two replicates. Over 97% of insertions occurred within annotated features at an average density of 9 unique insertions per kb (25). Despite opportunities for bottlenecking and bias during each growth, extraction, labeling and sequencing step, the number of reads mapped per annotated feature was reproducible with a Pearson’s coefficient of 0.98 between the two culture replicates (Fig. 1A). When two independent cultures were further grown with poised electrodes as the electron acceptor, then separately recovered, extracted, and labeled, the number of reads mapped per gene in each independent growth experiment was similarly reproducible with a Pearson’s coefficient of 0.98 (Fig. 1B).

**Figure 1.**
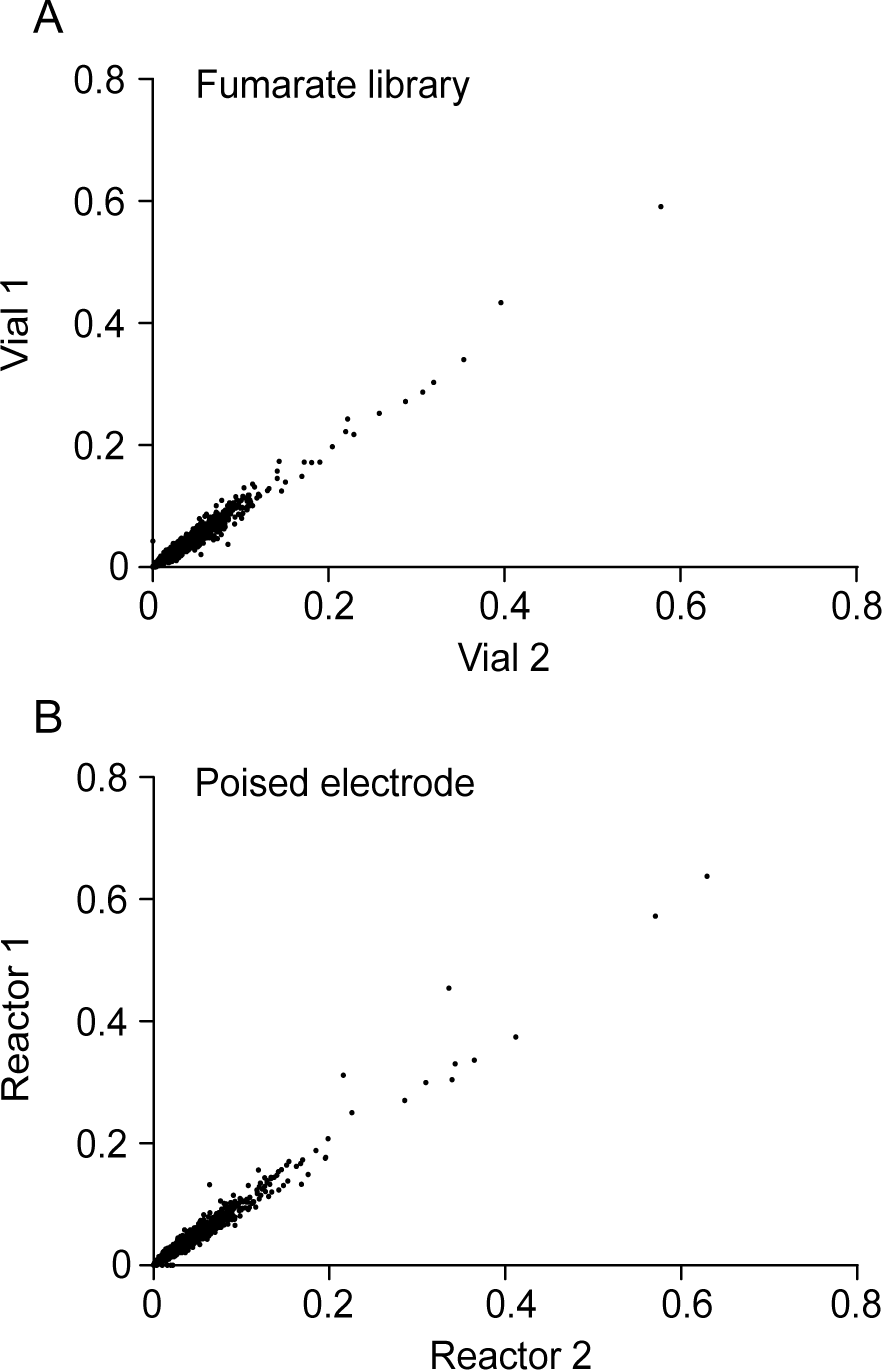
Tn-Seq reproducibility within library replicates and between experimental replicates in *Geobacter sulfurreducens*. (A) Comparison of two subsamples of the same fumarate-grown library. The number of reads mapped to a gene (normalized for read depth) is plotted against data for the same gene prepared and sequenced in parallel. (B) Comparison of two separate cultures inoculated on a poised electrode as terminal electron acceptor, recovered and sequenced separately.

### Essential genes for growth using fumarate in *G. sulfurreducens*

To predict genes important for growth with electrodes, genes required for growth in standard minimal medium with the electron acceptor fumarate were first determined, so they could be excluded from further analyses. Tn-Seq libraries were grown in liquid culture for 6 generations and sequenced to a depth of >40 M reads per culture replicate. At this depth, an average of 1,200 reads are obtained per insertion, and based on an average feature size in *G. sulfurreducens* of 0.95 kb/feature, 10,200 reads are expected to map to each feature in the annotated genome.

Previous constraint-based modeling predicted that a set of 140 central metabolic and biosynthetic genes should be essential for growth using fumarate as the sole electron acceptor (26). The majority (>90%) of these predicted essential genes had less than 300 mapped reads in our Tn-Seq analysis, compared to an expected 10,200 reads/gene, or they contained less than 4 insertions per kb (Fig. 2, Table S2). A small subset of genes previously predicted to be essential *in silico* averaged over 8,100 reads/feature and 10 insertions per kb, similar to other non-essential genome regions, and were considered to be non-essential under our growth conditions. Based on these results, annotated features with less than 300 reads mapped per feature or 4 insertion sites per kb were categorized as essential. Using both criteria allowed correction for small genes with few transposon insertion sites (TA dinucleotide) as well as larger genes which can support insertions between domains (27). According to this cutoff, over 170 genes previously coded as non-essential in the *in silico* model were essential in our minimal medium. Most of these encoded biosynthetic pathways for biotin, riboflavin, and cobamide, as our medium lacked vitamins. In addition, biosynthesis pathways for heme, fatty acids, and lipopolysaccharides, along with putative transporters for sulfate, acetate, copper, cobalt, and magnesium were essential. Complete tables comparing *in silico* predictions with Tn-Seq results are in Tables S3-S4.

**Figure 2.**
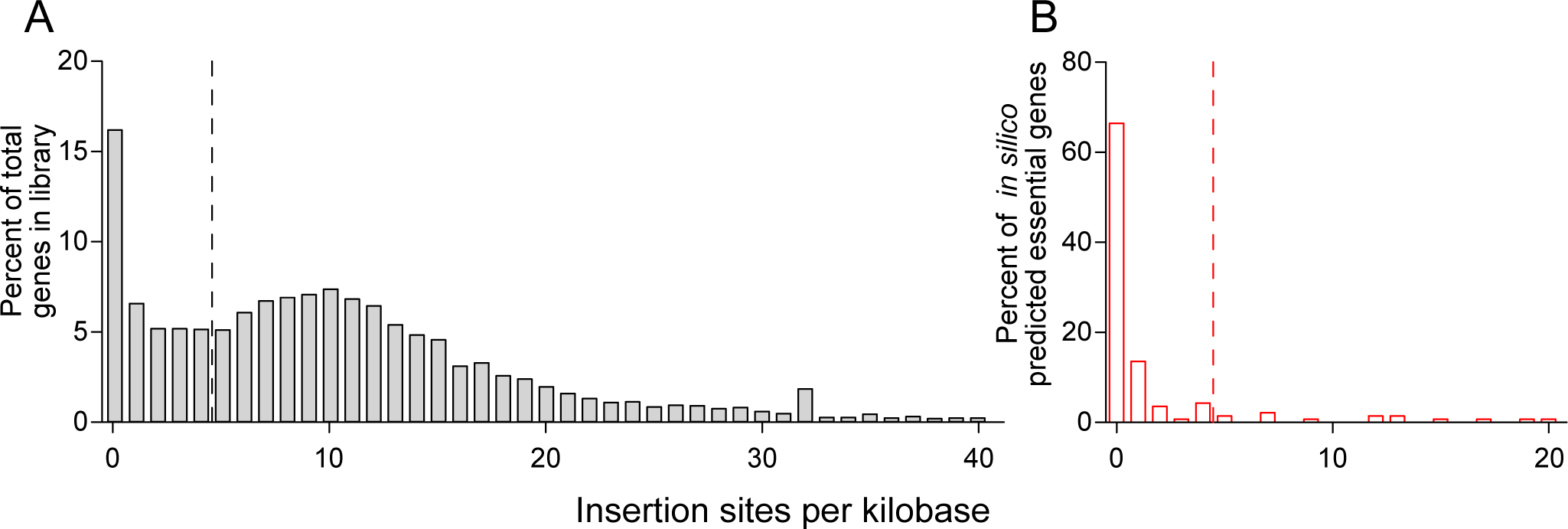
Estimation of essential genes based on low insertion densities, and use of *in silico* model data to verify essentiality predictions. (A) The frequency distribution for fumarate-grown cells follows a bimodal distribution centered around 10 insertions/kb, with strong enrichment below 4 insertions per kb (left of the dashed line). Genes with few insertions are predicted to be essential under these conditions. (B) Most genes labeled as essential in *in silico* modeling also contained less than 4 insertion sites per kb, and had less than 300 mapped reads/gene (26).

In total, almost 40% of annotated features, or 1,214 genome features out of 3,706, were identified as essential for growth with acetate as the electron donor and fumarate as the sole electron acceptor. A key difficulty in the original *G. sulfurreducens* annotation and *in silico* metabolic model was that genes encoding homologs or redundant pathways were not assigned function, as they could possibly complement each other (18, 19). Tn-Seq analysis clarified hundreds of such issues. For example, despite the presence of five annotated ferredoxins, only one was essential (GSU2708), one of three 2-oxoglutarate dehydrogenase complexes was essential (GSU1467 – GSU1470), one of three aconitase enzymes was essential (GSU1660), one of two fructose bisphosphate aldolases was essential (GSU1193), and two of three phosphoglycerate mutase enzymes were essential (GSU1818 and GSU3207). In total, Tn-seq resolved the essentiality of over 270 such examples where homologs were present but only specific genes were required under laboratory conditions (Table S5-S6).

Of special interest in *Geobacter* electron transfer is the mechanism of NADH oxidation and electron delivery to the quinone pool. This remains poorly understood, partially due to the fact that most *Geobacteraceae* encode two distinct complex I-NADH dehydrogenases (28). The complex I similar to that found in other Proteobacteria, recently classified as type “E” (GSU3429 – GSU3445) was not essential in *G. sulfurreducens*. However, a second, less well characterized complex I was essential. This NADH dehydrogenase does not fall into a major defined clade, and is most closely related to complex I sequences in *Chloroflexi* (GSU0338 – GSU0351) (28). A second essential gene cluster with bioenergetic implications was an integral membrane *b*-type cytochrome/FeS cluster/electron transfer flavoprotein. These genes (GSU2795 – GSU2797) encode all signature residues of confurcation/bifurcation complexes, suggesting high and low potential electrons (such as NADH + reduced Fd) may be combined for quinone pool reduction or proton translocation events during oxidation of acetate (29).

### Genes affecting electrode growth of *G. sulfurreducens*

Growth on an electrode surface in microbial fuel cells and electrochemical devices is hypothesized to be a complex phenotype that requires attachment, movement of electrons across membranes, and formation of between-cell conductivity to sustain respiration by cells not in contact with the surface (9, 10, 14, 30, 31). To test the entire process from attachment to biofilm growth, two parallel Tn-Seq libraries were grown for 6 generations on an electrode poised at -0.1 V vs. Standard Hydrogen Electrode (SHE), and the density of reads mapped per feature compared to parallel fumarate-grown cells. The electrode potential (-0.1 V vs. SHE) was chosen to mimic the redox potential of environmental Fe(III) oxides (32).

Expressing Tn-Seq read density data as Log2 ratios allows quick assessment of a phenotype. For example, a mutation preventing growth in a mutant results in twice as many wild type reads after one generation, or a Log2 ratio of -1. Using the exponential growth equation, genes with a Log_2_ score of -2 after 6 generations are equivalent to at least a 50% reduction in apparent doubling time. Using this criteria, nearly 50 genes encoding cytochromes, protein processing systems, sensing proteins, extracellular structures, polysaccharides, metabolic enzymes and hypothetical proteins were involved in electrode growth. Full data showing all read mapping densities and Log_2_ ratios for all features is in Table S2, and all raw read data needed to re-create this analysis is available in NCBI BioProject PRJNA290373. Direct links for downloading read mapping files and the annotated genome for viewing in IGV is in file S7.

Tn-Seq analysis confirmed a previous finding that CbcL, an inner membrane putative quinone oxioreductase containing both *b*- and *c*-type cytochromes (GSU0274) is crucial for electrode growth when electrodes are poised at -0.1 V vs. SHE (9, 10). Insertions in *cbcL* resulted in Log_2_ ratios corresponding to doubling times of 17.4 h, while a ~20 h doubling time was previously estimated for a pure culture Δ*cbcL* mutant (9). Also in agreement with previous work was the severe negative impact of insertions in genes involved in assembly and expression of the conductive Type VI pili. Insertions in the *pilQPOM* operon (GSU2028 – GSU2032), *pilC* gene (GSU1493) and *pilS* sensor kinase (GSU1494) produced similar defects, while insertions in other Type VI pili genes predicted weaker phenotypes (below the threshold in Table 2), but still causing at least a 25% decrease in apparent growth rate. Insertions in some pili genes, such as the *pilT4* gene (GSU1492), led to increased apparent abundance, consistent with recent reports that pilT mutants more rapidly colonize electrodes (33).

**Table 1.**
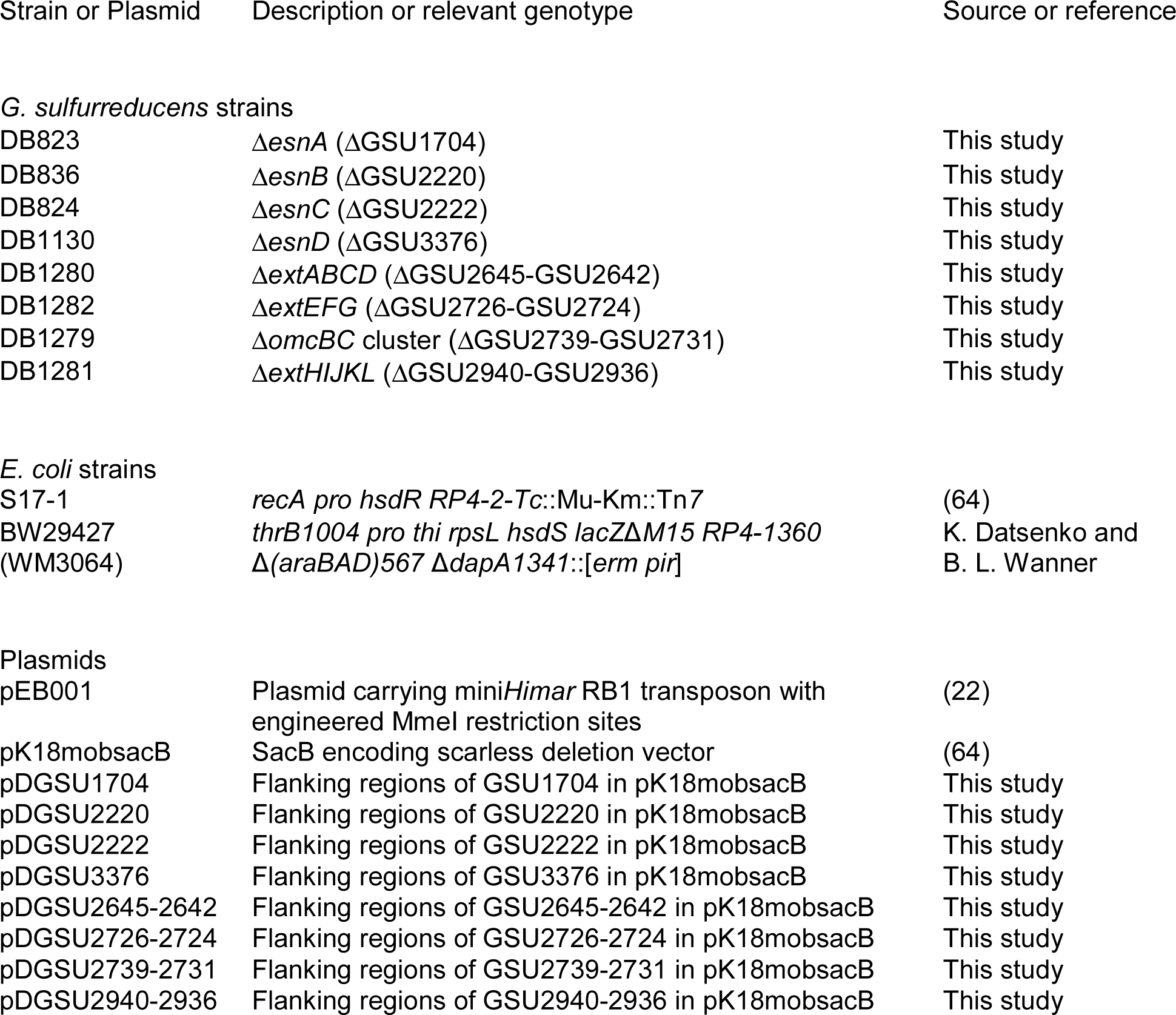
Strains and plasmids used in this work

| Strain or Plasmid | Description or relevant genotype | Source or reference |
| --- | --- | --- |
| *G. sulfurreducens* strains |  |  |
| DB823 | $\Delta esnA$ (ΔGSU1704) | This study |
| DB836 | $\Delta esnB$ (ΔGSU2220) | This study |
| DB824 | $\Delta esnC$ (ΔGSU2222) | This study |
| DB1130 | $\Delta esnD$ (ΔGSU3376) | This study |
| DB1280 | $\Delta extABCD$ (ΔGSU2645-GSU2642) | This study |
| DB1282 | $\Delta extEFG$ (ΔGSU2726-GSU2724) | This study |
| DB1279 | $\Delta omcBC$ cluster (ΔGSU2739-GSU2731) | This study |
| DB1281 | $\Delta extHIJKL$ (ΔGSU2940-GSU2936) | This study |
| *E. coli* strains |  |  |
| S17-1 | *recA pro hsdR RP4-2-Tc::Mu-Km::Tn7* | (64) |
| BW29427<br>(WM3064) | *thrB1004 pro thi rpsL hsdS lacZΔM15 RP4-1360*<br>$\Delta(araBAD)567 \Delta dapA1341::[erm pir]$ | K. Datsenko and<br>B. L. Wanner |
| Plasmids |  |  |
| pEB001 | Plasmid carrying mini*Himar* RB1 transposon with<br>engineered Mmel restriction sites | (22) |
| pK18mobsacB | SacB encoding scarless deletion vector | (64) |
| pDGSU1704 | Flanking regions of GSU1704 in pK18mobsacB | This study |
| pDGSU2220 | Flanking regions of GSU2220 in pK18mobsacB | This study |
| pDGSU2222 | Flanking regions of GSU2222 in pK18mobsacB | This study |
| pDGSU3376 | Flanking regions of GSU3376 in pK18mobsacB | This study |
| pDGSU2645-2642 | Flanking regions of GSU2645-2642 in pK18mobsacB | This study |
| pDGSU2726-2724 | Flanking regions of GSU2726-2724 in pK18mobsacB | This study |
| pDGSU2739-2731 | Flanking regions of GSU2739-2731 in pK18mobsacB | This study |
| pDGSU2940-2936 | Flanking regions of GSU2940-2936 in pK18mobsacB | This study |

**Table 2.**
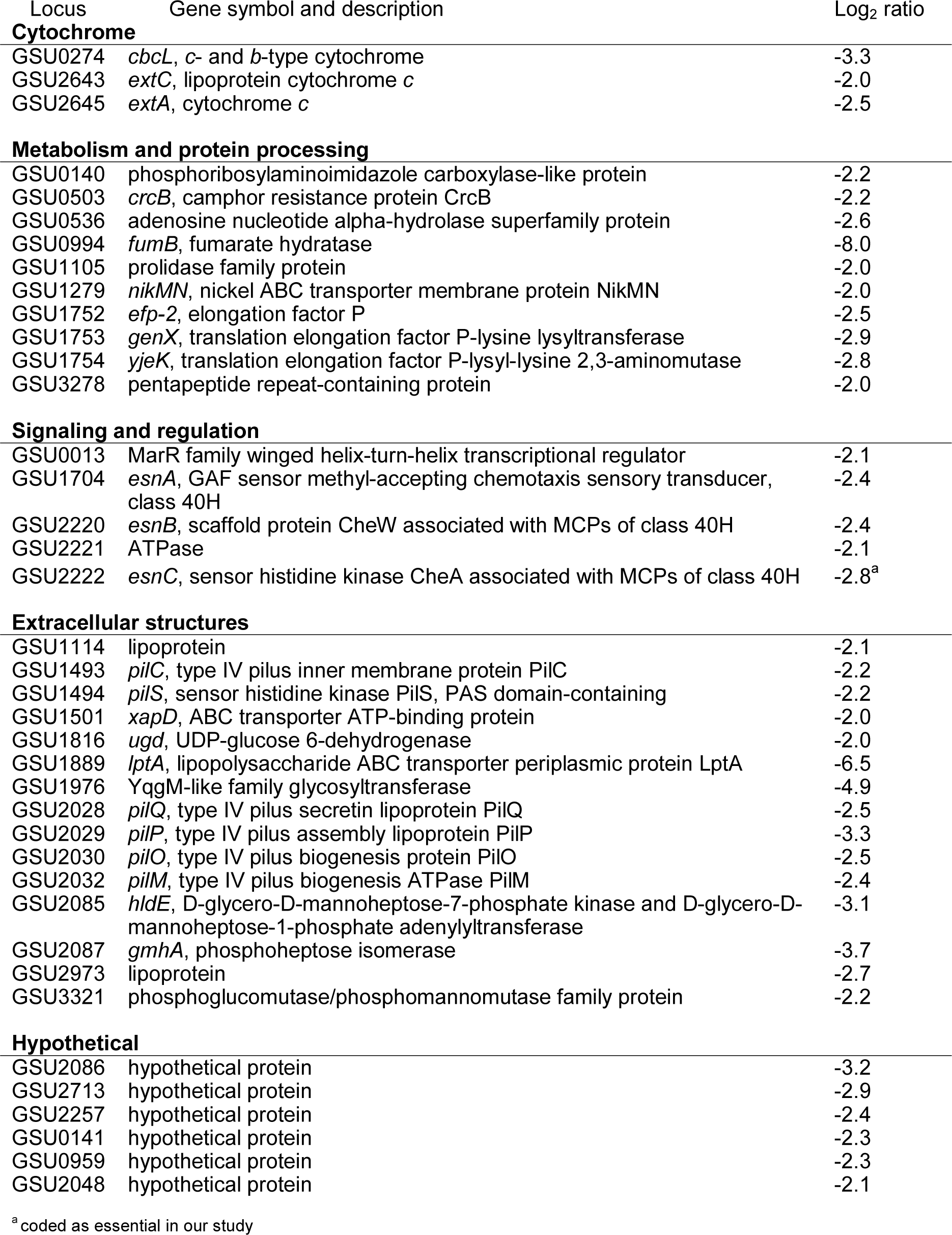
TnSeq mutations which after growth on an electrode showed a decrease in reads mapped by at least a Log_2_ ratio of -2, equivalent to a predicted 50% reduction in growth rate.

Surprisingly, with the exception of CbcL, no other cytochromes reported to play a role in electron transfer to metals appeared to be required for growth with an electrode in the Tn-Seq analysis. Instead, insertions in two uncharacterized multiheme *c*-type cytochromes did significantly affect electrode growth (GSU2643 and GSU2645). These cytochromes are part of a cluster containing two multiheme *c*-type cytochromes, a multiheme lipoprotein cytochrome, and a predicted outer membrane porin-like protein. As a similar arrangement of multiheme cytochromes, lipoprotein cytochromes, and putative β-barrel proteins encodes a “conduit” for electron transfer across the outer membrane in both *Geobacter* (the OmcB-based conduit) and *Shewanella* (the MtrCAB conduit), this region was targeted for further study.

The *G. sulfurreducens* genome encodes over 100 histidine kinases, response regulators and chemotaxis-like proteins, yet little is known about their role controlling phenotypes such as extracellular respiration (34). Tn-Seq identified one DNA-binding protein (GSU0013), and four chemotaxis proteins (GSU1704 and GSU2220 – GSU2222) as significant at the level of a 50% growth reduction. Only one diguanylate cyclase-response regulator protein was suggested to participate in electrode-dependent growth (GSU3376) above the level of a 25% reduction in growth. In model systems, methyl-accepting chemotaxis proteins such as GSU1704 undergo a conformational change to interact with CheW (GSU2220) and CheA (GSU2222) family proteins, triggering phosphorylation of GSU3376-like response regulators (35). Based on the hypothesis that GSU1704, GSU2220, GSU2222 and GSU3376 represented portions of MCP-CheA-CheW-response regulator systems involving the biofilm regulator c-di-GMP, these were also targeted for further study.

Some of the strongest phenotypes in apparent electrode growth were due to insertions in genes encoding sugar and polysaccharide synthesis. Five different sugar dehydrogenases, sugar transferases, and isomerases were identified as being required for electrode growth via Tn-Seq, and insertions within at least three different gene clusters involved in capsule, lipopolysaccharide and extracellular sugar synthesis produced significant defects. Also crucial were genes putatively involved in translocating sugars to the outer surface, such as a sugar ABC transporter previously shown to be essential for electrode growth (*xapD*, GSU1501), and an LPS ABC transporter protein (*lptA*, GSU1889). The defects created by these insertions further support a role for sugars in creating a properly charged or modified outer surface during *Geobacter* biofilm formation on electrodes (14, 16).

Finally, proteins involved in translation and processing of proline-rich proteins became important during electrode growth. For example, elongation factor P is essential for overcoming ribosome stalling during translation of proline-repeat sequences, and contributes to outer membrane integrity in *E. coli* (36, 37). Tn-Seq analysis predicted that one of two Ef-P homologs in *G. sulfurreducens*, (Efp-2, GSU1752), along with the Ef-P modifying Ef-P lysine-lysyltransferase (GSU1753) and Ef-P lysyl-lysine 2,3-aminomutase (GSU1754) became important during growth with electrodes. Insertions in a prolidase involved in cleaving peptides at proline residues (GSU 1105) and peptidylprolyl cis-trans isomerase (GSU2074) showed strong growth defects, further suggesting an unrecognized importance in the folding and processing of proline-rich proteins under electrode respiring conditions.

### Possible genes missed by the community aspect of Tn-Seq analysis

Deletion of the outer surface cytochrome gene *omcZ* (GSU2076) can significantly decrease electrode growth (38). While Tn-Seq insertions in genes within the *omcZ* operon had a negative impact (such as the peptidylprolyl cis-trans isomerase, GSU2074), insertions in *omcZ per se* were not identified as causing a defect in our library experiments (38, 39). Similarly, the pili-associated cytochrome OmcS is reported to be involved in electron transfer to electrodes, yet it was not identified in our analysis. Both of these cytochromes are verified to exist beyond the cell membrane, in the conductive matrix between cells (39, 40).

Under Tn-Seq conditions, enzymatic activities and proteins can be shared. A clear example of population-based complementation was evident in Tn-Seq data for fumarate hydratase, a TCA cycle gene predicted *in silico* to be absolutely essential (GSU0944, Table S3). Due to the fact that wild type *G. sulfurreducens* secretes malate when grown with fumarate as the electron acceptor (41), we hypothesized that fumarate hydratase was not required by the small subpopulation of fumarate hydratase mutants, as extracellular malate can rescue the gap in the TCA cycle. Consistent with this hypothesis, during growth with an electrode in the absence of fumarate, fumarate hydratase became one of the most essential genes in Tn-Seq analysis.

Our data suggest that extracellular cytochromes such as OmcZ secreted by wild type cells may similarly aid the occasional Δ*omcZ* mutant in the biofilm population during Tn-Seq experiments. This phenomenon would be similar to how outer membrane proteins shared between *Myxococcus* strains rescue motility mutants (42), and siderophores act as “public goods” for rare non-producing strains (43). In contrast to evidence that OmcS and OmcZ might be shared between cells, cellular machinery such as the Type IV pili, inner membrane cytochromes and outer membrane cytochromes remained essential even in biofilm conditions. Further Tn-Seq experiments may provide a mechanism to discover which *Geobacter* extracellular proteins can be shared for communal conductivity and exploited by ‘cheaters’ embedded in a conductive matrix, vs. which proteins catalyze key attachment or electron escape reactions so essential to each cell that they cannot be borrowed from neighbors.

### Cytochrome conduit deletion mutants affect electrode-but not Fe(III)-reduction

While Tn-Seq agreed with the importance of previously reported mechanisms such as pili, inner membrane cytochromes, and extracellular sugars, it identified many processes never highlighted in any mutant, proteomic, or transcriptional study. Based on their strong phenotypes under Tn-Seq conditions, we investigated the roles of two surprising classes of mutants, involving new outer membrane cytochromes and signaling proteins, after constructing scarless deletions of these key genes and gene clusters.

In Tn-Seq data, no effect was observed for mutations in one of the most studied outer membrane *c-*type cytochromes in *G. sulfurreducens*, OmcB. Part of a trans-outer membrane porin-cytochrome conduit capable of transferring electrons across the outer membrane, OmcB is encoded in a three-gene cytochrome/porin-like protein/lipoprotein cytochrome cluster (*ombB-omaB-omcB*, GSU2737–GSU2739) located next to a nearly identical tandem duplication containing another conduit (*ombC-omaC-omcC*, GSU2731–GSU2733). This duplication limits the impact of single mutations in any one gene. Thus, a scarless deletion of this entire genomic region, lacking all genes in the *omcB* and *omcC-*encoded conduits was constructed, and is referred to here as Δ*omcBC* (ΔGSU2739 – GSU2731). cytochrome conduit, that shares no homology with the OmcB or OmcC conduits, showed a strong phenotype in Tn-Seq data. This conduit was termed *extABCD* (extracellular electron transfer) and genes were removed to generate the Δ*extABCD* (ΔGSU2645-GSU2642) strain. In addition, deletion mutants of two other putative outer membrane conduit clusters, Δ*extEFG* (GSU2726 – GSU2724) and Δ*extHIJKL* (GSU2940 – GSU2936) were constructed as controls to compare with Tn-Seq results.

The Δ*extABCD* strain grew poorly when the electrode was the electron acceptor, never producing more than 50 µA/cm^2^ when grown under the same conditions used for Tn-Seq. In contrast, the Δ*omcBC* deletion strain showed no defect, and grew on the electrode similar to wild type (Fig. 3A). The other mutants lacking outer membrane conduits, Δ*extEFG* and Δ*extHIJKL*, also showed no defects on the electrode, demonstrating similar growth rates and final current densities as wild type. All mutants grew with wild type growth rates using fumarate as the electron acceptor, as predicted by Tn-Seq essentiality data (data not shown). While single-gene replacement mutants lacking the *omcB* gene will grow in microbial fuel cells (44), these results demonstrated that no part of the *ombB-omaB-omcB* and/or *ombC-omaC-omcC* conduit genes were required for wild type colonization and reduction of electrodes. Two other putative trans-outer membrane conduit gene clusters are also not essential for electron transfer to electrodes. In both Tn-Seq and in pure culture studies with reconstructed mutants, only the deletion of *extABCD* had any effect when electrodes were the electron acceptor.

**Figure 3.**
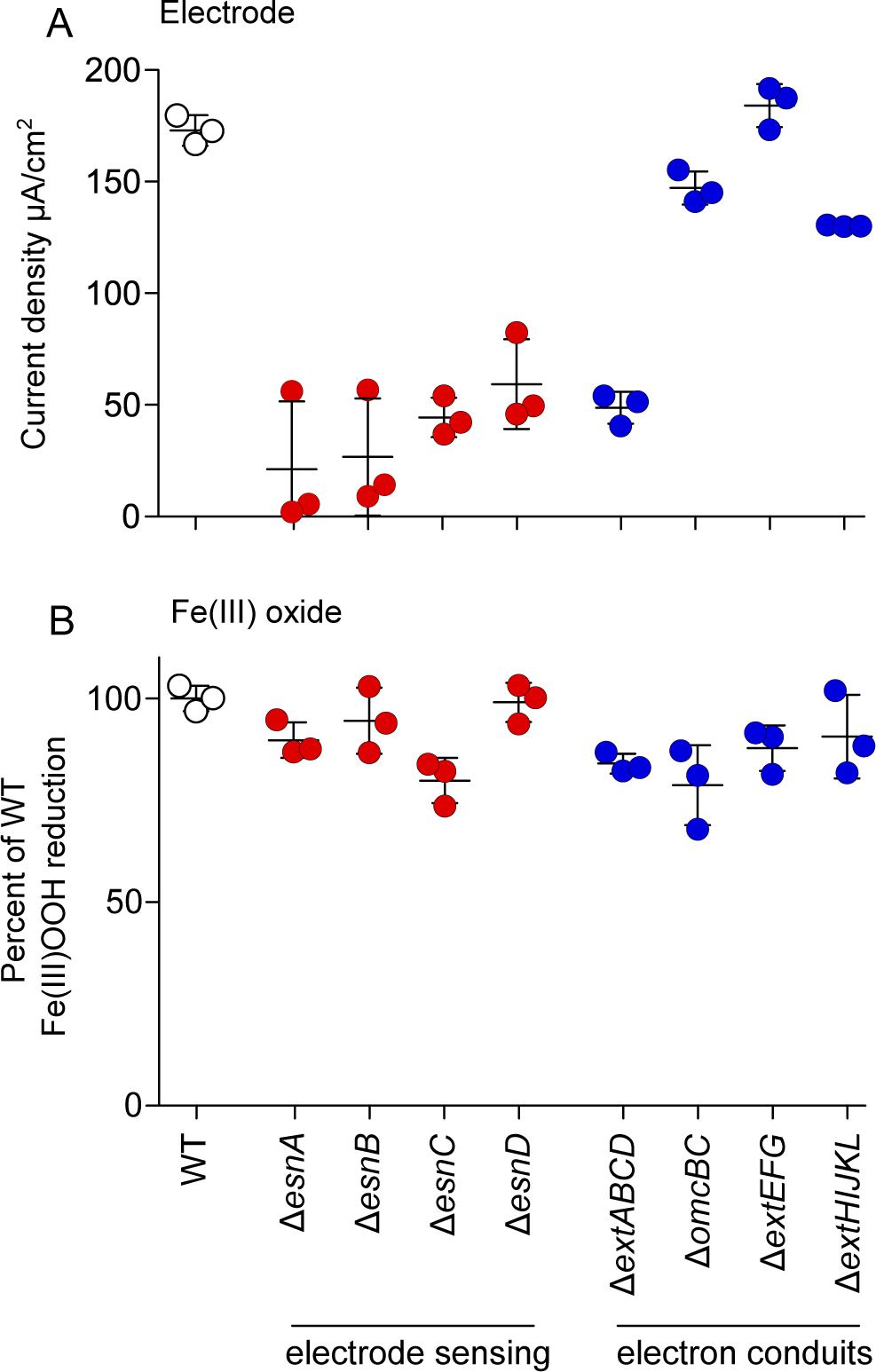
Genes essential for growth with electrodes are not required for Fe(III) reduction. Growth of scarless deletion mutants of chemosensory-like genes *esnA, esnB, esnC* and *esnD* and extracellular conduit clusters *extABCD, omcBC, extEFG* and *extHIJKL* with (A) insoluble Fe(III) oxides and (B) poised electrodes. The amount of Fe(III) reduced in 7 days was normalized to WT. Maximum current densities on electrodes poised at -0.1 V vs. SHE were recorded for all strains 80 hours after inoculation. All experiments were performed in triplicate.

In contrast to the strong electrode phenotype observed for the mutant lacking *extABCD*, reduction of insoluble Fe(III)-oxide by Δ*extABCD* was unaffected. Even though metal oxide particles are surfaces with similar redox potentials as the electrodes used in the Tn-Seq analysis, the *extABCD* gene cluster was not essential for reduction of metals (Fig. 3B). The phenotype of the Δ*extABCD* mutant suggests that the dominant pathway of electron transfer across the outer membrane will vary, based on what is being used as the extracellular electron acceptor.

### Chemosensory system deletion mutants also affect electrode-but not Fe(III)-reduction

A second class of genes revealed by Tn-Seq analysis were related to intracellular sensing and signaling systems. Based on their Tn-Seq phenotypes, these proteins were termed part of an <u>e</u>lectrode <u>s</u>ensing <u>n</u>etwork, and included; EsnA, a methyl-accepting chemotaxis protein (GSU1704), EnsB, a CheW-like chemotaxis scaffolding protein (GSU2220), EsnC, a CheA-like chemotaxis histidine kinase (GSU2222), and EsnD, a diguanylate cyclase (GSU3376). Scarless deletion mutants of each *esn* gene were constructed and tested for growth using an electrode as the electron acceptor.

All four *esn* deletion mutants displayed defects in growth with an electrode even more severe than the Log_2_ ratios observed in Tn-Seq data (Fig. 3A and Table 2). However, when these same four *esn* mutants were grown with Fe(III)-oxide as the electron acceptor, Fe(III) reduction was similar to wild type (Fig. 3B). All mutants also grew at wild type growth rates with fumarate as the electron acceptor. This phenotype, where a mutant performed poorly with an electrode surface but still reduced metal oxide particles, agreed with the hypothesis that cells interact with these two solid-phase electron acceptors via different molecular mechanisms.

Further work will be required to determine if the similar phenotypes displayed by *esn-*encoded proteins is due to direct physical interactions or common signaling molecules. No annotated MCP genes in the *G. sulfurreducens* genome are co-localized in operons with a complete set of CheA-CheW genes, and no protein-protein interaction data is available to predict the downstream target of the EsnC CheA-like kinase. The EsnA methyl-accepting chemotaxis protein has no periplasmic sensing domain to suggest an external input for this system, but EsnA does contain a cytoplasmic GAF domain typically involved in binding small molecules such as cyclic nucleotides, and EsnD is a demonstrated cyclic-di-GMP producing response regulator (45). One hypothesis is that EsnABC together regulates and/or responds to levels of EsnD-produced cyclic-di-GMP, driving and reinforcing a switch to the conductive biofilm mode of growth (46). Whatever the ultimate signal is for electrode colonization, it does not appear to be involved in growth with metals such as Fe(III).

### Conclusions and outlook

The ability of Tn-Seq to survey mutants across the entire genome in a single growth experiment is especially useful when no high-throughput assay is available for the phenotype under study. In the case of growth on electrodes, testing 33,000 individual mutants in replicate electrode reactors would require our laboratory to dedicate its bank of 45 electrochemical cells to testing mutants nonstop for over 28 years. In a fraction of that time, saturation mutagenesis described over 1,200 genes essential for growth under laboratory conditions, resolved hundreds of annotation issues, and brought into focus over 50 genes vital during electrode growth conditions.

By asking what genes are required for growth, Tn-Seq generates data different from expression-based approaches which operate under the hypothesis that important genes will be strongly regulated in response to an electrode. Transcriptional data did lead to discovery of the key multiheme cytochrome OmcZ in *Geobacter*, and supported early hypotheses suggesting a role for pili. However, microarrays also found strong up-regulation of many genes which failed to show importance in follow-up mutant studies, and failed to show phenotypes in these Tn-Seq results (38). Examples include; the OmcB and OmcC cytochromes and multiple hypothetical cytochromes, amino acid transporters and hydrogenases, terminal oxidases involved in oxygen reduction, and heavy metal exporters. In general, we found little overlap between genes predicted as important to growth on electrodes via Tn-Seq and those up-or down-regulated in expression or proteomic studies on electrodes (31, 38, 47). One possibility is crucial regulatory or biosynthetic elements have low basal expression levels. Expression or abundance-based studies could have reduced sensitivity if biofilms are heterogeneous, or if expression varies with distance from the electrode (48). In the case of cytochromes, the *imcH* and *cbcL-*encoded cytochromes involved in electron transport are now known to be expressed even when cells are grown with fumarate, preventing their earlier discovery via differential expression (9, 10). It also remains possible that electrodes, being relatively unnatural substrates, trigger additional transcriptional changes or de-repress genes that have little to do with electrode respiration.

Tn-Seq data provided genome-scale information that agreed with prior *in silico* predictions and knockouts during fumarate reduction (26, 49). However, much less data are available for electrodes in terms of what genes are essential, and as with any large-scale survey, reconstruction of mutants is needed to corroborate findings. Using scarless mutants lacking genes and whole gene clusters, we were able to confirm that individual chemosensory components of *esnABCD* and the entire cytochrome conduit *extABCD* were essential for growth when the electrode acted as the electron acceptor. As two other outer membrane conduits, based on OmcB and OmcC, are well-studied and known to be involved in electron transfer to metals, we also constructed new mutants lacking these non-homologous cytochrome clusters, along with two other unstudied putative outer membrane conduits. Surprisingly, only removal of the *extABCD* conduit gene cluster affected electrode growth, as predicted by Tn-Seq.

When Fe(III) was the electron acceptor, none of the cytochrome conduit mutants with electrode phenotypes, and none of the mutants lacking methyl-accepting chemotaxis proteins or GGDEF-domain response regulators, had significant defects (Fig. 3B). These data support a model that *sulfurreducens* possesses separate mechanisms for reduction of electrodes compared to the reduction of metals that involves a distinct set of cytochromes for crossing the outer membrane.

In addition, separate cyclic dinucleotide-dependent regulatory systems may be used for recognition and respiration of electrodes compared to metals.

A recent comparison of seven *Geobacter* species noted a poor correlation between electricity production and Fe(III) reduction rates (50). In contrast, strains capable of high rates of electricity production were also capable of interspecies electron transfer to methanogens. This correlation led to the hypothesis that the conductive network accessible to electrodes evolved to support direct electron transfer between *Geobacter* and electron-accepting organisms such as methanogens. Evidence for this hypothesis lies also in the use of anaerobic digesters as the most common source of high current-producing *Geobacter* enrichments (51), compared to Fe(III)-rich environments (50, 52). A syntrophic partner may be much like an electrode; it can accept electrons indefinitely, but requires commitment to an attached biofilm lifestyle and the expense of building a conductive extracellular space (6, 53). In contrast, small metal oxide particles represent a temporary and more variable electron acceptor, where cells cannot attach permanently or grow into thick biofilms that entrap and re-use extracellular proteins (54).

While naturally-occurring *Geobacter* strains produce >1 mA/cm^2^ of electrical current, and this rate already rivals the best artificial enzyme-functionalized electrodes (55), use of microbial electrochemistry as an energy source or biocatalyst is estimated to require at least a 10-fold increase in current density in order to be profitable (56–58). Achieving these gains, and engineering these abilities into industrial bacteria, requires identification and enhancement of core components specific to electricity production. While the molecular basis for this extracellular respiration is becoming more clear, how organisms biochemically distinguish between insoluble metals vs. microbial or electrode-based acceptors remains a key challenge.

## Materials and Methods

**Growth and medium conditions.** All strains and plasmids used in this study are listed in Table 1. *G. sulfurreducens* strains and mutants were grown from single colony picks streaked from lab DMSO stocks in anoxic basal medium as described (59). For routine growth, basal medium with acetate (20 mM) as the electron donor and fumarate (40 mM) as the electron acceptor was used. Agar (1.5%) was added to the acetate-fumarate medium when culturing for clonal isolates on semisolid surface in an H_2_:CO_2_:N_2_ (5:20:75) atmosphere in a vinyl anaerobic chamber (Coy) or an anaerobic workstation 500 (Don Whitley). When electrodes were used as the electron acceptor, fumarate was replaced with 50 mM NaCl to maintain a similar ionic strength. When Fe(III)-oxide was used as the electron acceptor, fumarate was omitted and a non-chelated mineral mix was used (in which all components of the chelated mix were dissolved in a small volume of 1N HCl without NTA). In all cases, the pH of the medium was adjusted to 6.8, buffered with 2 g/L NaHCO_3_ and purged with N_2_:CO_2_ gas (80:20) passed over a heated copper column to remove trace oxygen.

Three-electrode bioreactors were assembled as previously described (60). Briefly, graphite electrodes were polished using 1,500 grit wet/dry sandpaper and attached to platinum wire to serve as the working electrode. A bare platinum wire was the counter electrode. The potential of the working electrode was maintained at -0.10 V vs. standard hydrogen electrode (SHE) using a saturated calomel reference electrode and a VMP3 multichannel potentiostat (Biologic). Reactor headspace was degassed using 80:20 N_2_:CO_2_ prior to inoculation. Current was measured as an average over two minutes. For all pure culture mutant experiments, a 25% inoculum with cultures reaching acceptor limitation (OD_600_=0.50-0.55) was used. The total volume of each reactor was 15 ml, and the working electrode surface area was 3 cm^2^. To assay Fe(III) oxide reduction in *G. sulfurreducens*, late exponential growth phase cultures (OD_600_ = 0.5-0.55) grown using acetate-fumarate were used to inoculate 1:100 in minimal medium containing 20 mM acetate as the electron donor and 55 mM freshly precipitated β-FeO(OH) as the sole electron acceptor. A small sample of the medium was removed at regular intervals and dissolved in 0.5 N HCl for at least 24 hours in the dark. The acid extractable Fe(II) was measured using a modified FerroZine assay (59).

*Escherichia coli* was cultivated in lysogeny broth supplemented with 0.3 M 2,3-diaminopimelic acid and 50 µg/ml kanamycin when needed.

### Tn-Seq library construction

Five ml of mid-log (OD_600_ ~0.35) *G. sulfurreducens* culture grown on acetate-fumarate was combined with 5 ml of an overnight culture of *E. coli* conjugative donor strain BW29427 (WM3064) carrying the transposon plasmid pEB001 was applied to a nitrocellulose filter (0.4 μm pore size) using a vacuum. Plasmid pEB001 contains the *mariner* derivative *Himar1* transposon with MmeI recognition sites on both sides of the inverted repeats that flank a kanamycin resistance cassette. *G. sulfurreducens* recipient and *E. coli* donor mixture was washed on the filter with 3 volumes of basal medium before transferring the filter with the cell mixture to an agar plate containing acetate-fumarate and incubated in a Coy chamber at 30°C for 4 hours. The cell mixture was washed off the filter disc with 1 ml of basal medium supplemented with 200 µg/ml kanamycin. To select for *G. sulfurreducens* transposon mutants, dilutions were plated on large 22 x 22 cm square agar plates with acetate-fumarate medium supplemented with kanamycin and incubated at 30°C in the anoxic chamber until visible colonies formed (in six days). Approximately 50,000 colonies were pooled in 10 ml of acetate-fumarate medium and grown for 4 hours before freezing 1 ml aliquots in 10% DMSO.

### Tn-Seq experiment

For each experiment, a frozen library aliquot was used to inoculate 100 ml of acetate-fumarate medium. When cultures reached exponential growth phase (OD_600_ ~0.35), 5 ml of this culture was inoculated into fresh 100 ml acetate-fumarate medium. This parent culture was grown to early stationary phase (OD_600_ ~0.5) and used to initiate both electrode and fumarate control experiments. Five ml of the parent culture was inoculated into 100 ml of acetate-fumarate medium and allowed to grow for 6 generations before harvesting. At the same time, a 7.5 ml of the parent culture was inoculated into the 3-electrode bioreactor containing 7.5 ml of medium, where two 3 cm^2^ working electrodes were present to support biofilm growth. *G. sulfurreducens* electrode biofilms were harvested by moving reactors to an anaerobic chamber after 6 generations of growth using electrodes poised at -0.1 V SHE, with the number of generations estimated based on the doubling time (10 h) at this redox potential. The entire electrode plus attached biofilm was placed in 100 ml of acetate-fumarate minimal medium, vortexed to liberate cells, and incubated at 30°C for 6 generations before harvesting. This outgrowth was necessary to produce adequate free biomass for DNA extraction and sequencing, and is why cells grown with fumarate for an additional 6 generations were used as a parallel control. Despite this additional outgrowth step, read densities and phenotypes between replicate experiments were highly repeatable (see Fig 1B).

Genomic DNA from 40 ml of cells was isolated using the Wizard genomic DNA purification kit (Promega). The protocol for preparing the DNA library for Illumina sequencing is outlined with modifications (61). Six µg of gDNA was digested with MmeI (New England Biolabs) for 2 hours at 37°C. To this reaction, Antarctic phosphatase (New England Biolabs) was added and incubated at 37°C for an additional hour. Enzymes were inactivated at 65°C for 15 minutes, followed by phenol/chloroform/isoamyl acetate (25:24:1) extraction and ethanol precipitation at - 20°C overnight. MmeI digested gDNA and Illumina barcoded adaptor with two random base pair overhangs were ligated using T4 DNA ligase (Epicentre) for 1 hour at 25°C. The transposon with 20 bp of genomic DNA sequence junction and ligated adaptor was amplified using Phusion High GC master mix (New England Biolabs) using primers (P1 M6 MmeI and Gex PCR Primer 2) that anneal to the inverted repeat of the transposon and the ligated Illumina adaptor. The PCR reaction was terminated during linear amplification (24). The 120 bp product was gel purified and saved at -20°C. After Sanger sequencing verification using primer Gex PCR Primer 2 for the presence of the unique barcode, the PCR product was sequenced using Illumina (HiSeq 2500 Rapid chemistry single read 50 bp). Three samples with unique barcodes were mixed in a single Illumina lane generating ~30 M quality-passing reads per sample. All sequencing was performed by the University of Minnesota Genomics Center. Primers and barcoded adaptors used in the construction of the Tn-Seq library are referenced here (61).

### Mapping of transposon insertions

For each Tn-Seq library, raw sequences were de-multiplexed and extracted according to the unique barcode. The barcode and transposon sequences were trimmed, keeping only genome sequences that were at least 16 bp in length. Reads were then aligned to our *G. sulfurreducens* reference genome (59) using Bowtie (62) (Version 1.1.2) with no mismatches allowed, and discarding reads that could map in more than one location. In a typical experiment, more that 90% of reads were mapped to unique sites. After subtracting TA insertion sites in genes found to be essential under our conditions, the *G. sulfurreducens* genome contains about 55,000 sites where the *mariner* transposon could insert and produce viable mutants. As we also discounted insertions in the first 5% of a feature, as these often do not produce a knockout phenotype (27), libraries recovered insertions in over 64% of available genomic TA sites that could produce viable mutants, providing an average of nearly 10 independent mutants per gene feature.

Approximately 2.7% of the insertional reads mapped to more than one location in the *G. sulfurreducens* genome due to redundancies. Of interest to our analysis, the *omcB* (GSU2738 – GSU2737) and *omcC* (GSU2735 – GSU2731) gene clusters are tandemly duplicated in the genome. Two genes in each cluster are 99-100% identical (GSU2739-8 and GSU2733-2) at the nucleotide level, preventing unique readsmapping to these genes, while the 12-heme cytochrome genes *omcB* (GSU2737) and *omcC* (GSU2731) share 87% DNA sequence identity and contained a few characteristic sites. When the 20 bp MmeI-generated genomic DNA reads mapped ambiguously to more than one site, they were excluded from analysis, and genes containing such sites were flagged to account for possible “low insertion density” in the gene that would wrongly code it as essential. Similar issues limited the amount of useful data for the small triheme cytochromes encoded in *ppcA-E* because there are few unique 20 bp regions in these small genes adjacent to TA sites. However, most ambiguous reads mapped to the 35 transposable elements and duplicated ribosomal RNA genes. All ambiguous mappings were excluded from the essentiality analysis, but were included for total read normalization between barcoded libraries.

To estimate the severity of a phenotype, the effective change in doubling times for Tn-Seq mutants was calculated. The exponential growth equation was applied using the WT doubling time of 10 hours for the duration of electrode growth (72 hours), comparing the reads in a gene between electrode and fumarate conditions where x is the apparent electrode growth doubling time (hours). According to the conditions of the experiment, a Log_2_ ration of -2 after 72 hours implies a mutant with a doubling time of ~15 hours.

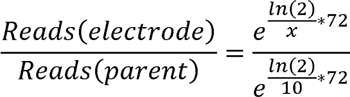

All of the Tn-Seq library sequence manipulations were performed in a Galaxy server hosted on the Minnesota Supercomputing Institute at the University of Minnesota (63).

### Scarless gene deletion in *G. sulfurreducens*

Roughly 1 kb flanking the targeted gene(s) of interest were cloned into the *sacB* encoding pK18mobsacB plasmid. To prevent disruption of the flanking genes, about 30 bp of the target gene sequence coding for a small peptide was cloned within the flanking region. The *sacB* plasmid was transformed into *E. coli* conjugative donor strain S17-1 to conjugate into *G. sulfurreducens* recipient. One ml of fully grown *G. sulfurreducens* acetate-fumarate culture was pelleted on top of 1 ml of S17-1 culture carrying the *sacB* plasmid, mixed on top of a 0.22 µm filter resting on acetate-fumarate agar plates in an anaerobic chamber and incubated for 4 hours before streaking the mixture onto acetate-fumarate plates with 200 µg/ml kanamycin. This procedure selected *G. sulfurreducens* culture with pK18mobsacB integrated into either flanking region of the gene since the plasmid cannot replicate in *G. sulfurreducens*. Scarless gene deletion mutant was selected on acetate-fumarate plates containing 10% sucrose and confirmed using PCR with primers flanking the deletion site (59). The primers used to clone the flanking regions into pK18mobsacB and flanking primers to confirm gene deletions are listed in Table S1.

### Tn-Seq raw sequencing data

Illumina sequence data trimmed to remove transposon sequences and barcode sequences, containing only genomic DNA (.fastq) from each condition are deposited in NCBI short read archives with the following accession number: SRX2199236 (fumarate outgrowth, used for determining essentiality), SRX2199234 (parent library, used as the reference to compare between fumarate and electrode conditions), and SRX2199233 (concatenated with fastq headers HJHKHADXX and HJF5FADXX for electrode outgrowth replicates). Mapping files (.bam) for the fumarate outgrowth dataset used to determine essentiality is deposited as SRX2199235. The reference *G. sulfurreducens* genome used for all mapping was re-sequenced and deposited as SRX1101230. Instructions to view read mapping against our reference genome using IGV is available in supplemental file S7.

## Acknowledgements

We thank M. Mehta for initiating the construction and analysis of preliminary Tn-Seq experiments in *G. sulfurreducens.* We thank J. Badalamenti for sequence analysis assistance and data management.

